# Stem cell mechanoadaptation Part B - Microtubule stabilization and substrate compliance effects on cytoskeletal remodeling

**DOI:** 10.1101/2024.07.28.605537

**Authors:** Vina D. L. Putra, Kristopher A. Kilian, Melissa L. Knothe Tate

**Affiliations:** School of Chemistry and School of Materials Science & Engineering, University of New South Wales, Sydney, NSW, Australia.; Blue Mountains World Interdisciplinary Innovation Institute (bmwi³), Blue Mountains, NSW, Australia

**Author notes:** Kristopher A. Kilian^1^, Melissa L. Knothe Tate^2^ **Email:**. **Author Contributions:** The studies were conceived of, and experiments designed by all named authors (VLDP, KAK, MLKT). The experimental studies were carried out and analyzed by VLDP under mentorship of KAK and MLKT. The data and analysis were interpreted by all named authors (VLDP, KAK, MLKT). VLDP wrote the manuscript which was discussed and revised and edited by all named authors (VLDP, KAK, MLKT). **Competing Interest Statement:** MLKT has co-founded start-up companies to commercialize the intellectual property she and her collaborators developed, and patent protected over the past decades. The topic of the current manuscript is fundamental in nature and not directly related to MLKT’s innovation translation and commercialization projects. **Classification:** Biological Sciences. Cell Biology, Biophysics.

**Keywords:** **C**ytoskeleton, Development, Mechanoadaptation, Stiffness

## Abstract

Stem cells adapt to their local mechanical environment by rearranging their cytoskeleton, which underpins the evolution of their shape and fate, as well as the emergence of tissue structure and function. Here we report on the second part of a two-part experimental series to elucidate spatiotemporal cytoskeletal remodeling and resulting changes in morphology and mechanical properties of cells, their nuclei, akin to mechanical testing of the most basic living and adapting unit of life, *in situ* in model tissue templates. We probed the native and PAX-exposed (inhibiting cytoskeleton tubulin depolymerization) stem cells’ cytoskeletal adaptation capacity on substrates of different compliance (exerting local tension on cells) and in combination with exposure to local compression effected with increased target seeding densities (5000 cells/cm^2^ - Low Density, LD; 15,000 cells/cm^2^, High Density, HD).

On 10 and 100 kPa gels, cells seeded at both LD and cells proliferated to HD exhibited bulk moduli that nearly matched those of their respective substrates, hence exhibiting a greater increase in Young’s Modulus after microtubule stabilization than cells cultured on glass. Culture on compliant substrates also reduced the PAX-mediated F-actin and microtubule concentration increase. On gels, F-actin alignment decreased as more randomly oriented, short actin crosslinks were observed, representing emergent adaptation to the compliant substrate, mediated through myosin II contractility.

We conclude that stem cell adaptation to compliant substrates facilitates the accommodation of larger loads from the PAX-stabilized polymerizing microtubule, which in turn exerts a larger effect in determining cells’ capacity to stiffen and remodel the cytoskeleton. Taken as a whole, these studies establish correlations between cytoskeleton and physical and mechanical parameters of stem cells that progress our understanding of the dynamic cytoskeleton, as well as shape changes in cells and their nuclei, culminating in emergent tissue development and healing.

**Significance Statement:** Stem cells adapt to their dynamic environment by means of cytoskeleton rearrangements - underpinning the emergence of tissue structure-function relationship; this represents a current gap in knowledge that needs to be addressed, to better target tissue neogenesis and healing in context of regenerative medicine. We introduced compression via increasing seeding density and tension via compliant substrates to create tissue templates, while stabilizing microtubules. We found that mechanical and biophysical cues exert a greater effect in modulating cytoskeletal adaptation than exogenous chemical agents targeting the cytoskeleton, thus counterbalancing the concentration-dependent effect on cell physical and mechanical properties. We further found that stem cells with stabilized microtubules are sensitive to a range of substrate stiffness and seeding density that allowed cells or multicellular constructs to broaden their capacity to adapt their mechanical properties.

## Introduction

Across the time and length scales of tissue development, stem cells adapt to the dynamic biomechanical and biophysical cues that determine the mechanical properties of their niche or tissue habitat^1,2^. Over time stem cells proliferate and grow, forming multicellular constructs with higher order architectures enabling more specialized functions. This process is exemplified *in situ* and *in vivo* by the process of tissue template (*Anlage*) development, *e.g.* formation of the condensing mesenchyme as the initial step in musculoskeletal development^2,3^.

Previously published work demonstrated the capacity of physiological stresses, *i.e.* intrinsic to growth, development, and maintenance of tissues^4,5^, to modulate cell and nuclear morphometries which tie closely to evolving cytoskeletal architectures, e.g. flattening of the nucleus via formation of an actin cap over and around the nucleus^6^. Such force-mediated modulation of cell and nuclear morphology has been shown to alter baseline gene expression of early mesenchymal condensation markers characteristic of emergent skeletogenesis as well as the expression of F-actin and tubulin filaments^7^. Following on our studies probing stem cell mechanoadaptation using exogenous modulators of cytoskeletal remodeling and local compression induced by seeding at increasing densities (see [Part A]), here (Part B) we examine effects of substrate compliance (or respective stiffness) on cell and nuclear morphology^8^,cytoskeletal remodeling^7^, and emergent lineage determination^9^.

We modulated the cells’ capacity to regulate stiffness and generate forces via exogenous exposure to Paclitaxel (PAX, which inhibits microtubule depolymerization), using methods described in the first part of these studies [Part A]. We hypothesized that the compliance of the substrate, together with local compression experienced by cells, and exposure to PAX, individually and together influence the spatiotemporal remodeling of the cytoskeleton. To that end we measured the spatial distribution of the cytoskeleton, changes in cell volume, shape and stiffness, and nucleus volume and shape over time periods relevant for early developmental processes (24h, 48h, 72h). By quantifying the time and PAX concentration-dependent changes in cell viability, proliferation, and cytoskeletal reorganization, we elucidated the cells’ mechanoadaptation capacity in contexts mimicking those which occur during early stages of tissue development.

## Results

### Substrate compliance modulates the cell volume increases and shape changes mediated by microtubule stabilization

To understand mechanoadaptation, as cells are subjected to as well as sense and counteract boundary forces, *e.g.* from the underlying substrate and/or surrounding cells, we introduced a range of polyacrylamide hydrogel substrates more compliant than glass, from stiffest gels (100 kPa and 10 kPa gels) to softest gels as compliant as cells themselves (1 kPa gels). To test the hypothesis that changes in stiffness at cell boundaries (substrate compliance) as well as intrinsic cell stiffness and size modulate force balances driving mechanoadaptation, we measured cell volume and corresponding actin filament alignment and cell stiffness, in association with culture on substrates of increasing compliance (decreasing resistance at cell boundaries) and PAX exposure (inhibiting tubulin depolymerization, increasing cell stiffness and volume).

### Effect of more compliant substrates on cell morphology

Substrate compliance exerts a profound influence on the local environment and subsequent adaptation of cell behavior (structure – function). Cell spreading was similar on the 10 and 100 kPa gel substrates; however, on the softest 1 kPa gel, most cells showed reduced spreading, and exhibited comparatively rounder morphology (Fig. 1A). Cells seeded on the stiffest, *i.e.* glass, substrate showed greater changes in morphology across observation timepoints than those seeded on more compliant gel substrates, where significant and grossly observable changes were observed first at the 72h timepoint (Fig. 1B). The cell volume increase associated with 100 nM PAX exposure shows comparable magnitude across the gel stiffnesses. However, with lower PAX concentrations on softer gels, cell volume remained small. As a result, cell shape measures (SA/V) were also generally smaller, indicative of rounder cells, for cells seeded on gels compared to those seeded on glass (Fig. 1C).

**Figure 1.**
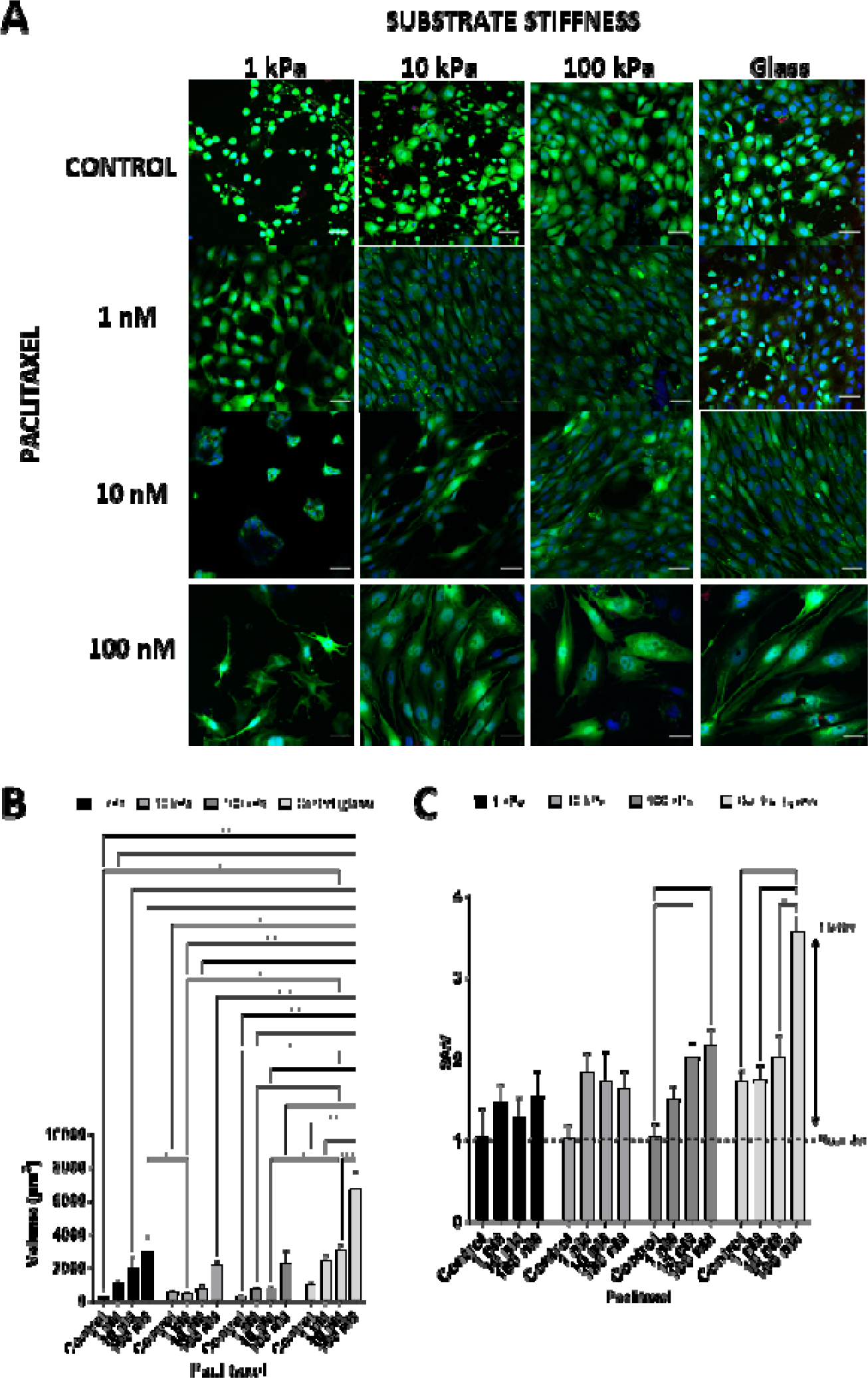
Compliant substrates modulate PAX concentration-dependent cell volume increases. (A) After 72h of PAX exposure, cells seeded on compliant gel substrates (scale bar 50 μm) show smaller volume and smaller increases in volume with increasing PAX concentration (B). With increasing substrate stiffness, cells exhibit average larger SA/V and the largest increase on glass (C) than those seeded gels due to their smaller volume.

Subtle changes in cell morphology (Fig. S1A), volume (Fig. S1B), shape (Fig. S1C) and spreading were observed as cells were adapting to softer substrates than to glass at 24h, although significant changes observed only in cells seeded on glass substrates. At 48h (Fig. S1D), PAX concentration-dependent cell volume increases were observed with increasing substrate stiffness (Fig.S1E), as was cell flattening (Fig.S1F).

### PAX-induced changes in cell volume and actin alignment regulate stem cell rigidity sensing that is enhanced in cells proliferating to high density

#### Effect of more compliant substrates on cytoskeleton filament concentration and spatial distribution

To study how cell volume adaptation on soft substrates translates to changes in cytoskeletal structure (Fig. 2A) we measured the spatial distribution and alignment of cytoskeleton in cells exposed to increasing concentrations of PAX and cultured on the 1, 10 and 100 kPa gels after 72h. Concentration-dependent cytoskeletal remodeling was observable as an increase in the amount both actin (Fig.2B) as well as microtubule (Fig. 2C) per cell with increasing concentration of PAX, for all groups, **except** cells exposed to 1 and 10 nM (lower concentrations) PAX and seeded on softest gel substrates (1 and 10 kPa). When we segmented actin and microtubules into the apical (Fig. 2D, E) and basal (Fig. 2F, G) regions of cells exposed to PAX and cultured on gel substrates, concentration-dependent differences nearly disappeared across all substrate compliances. Hence, early (over 72h in culture) mechanoadaptation of stem cells to substrates of different stiffnesses and in conjunction with PAX exposure appears to involve bulk increases in cytoskeletal proteins F-actin and tubulin without overt evidence of emergent cytoskeletal polarity at 72h.

**Figure 2.**
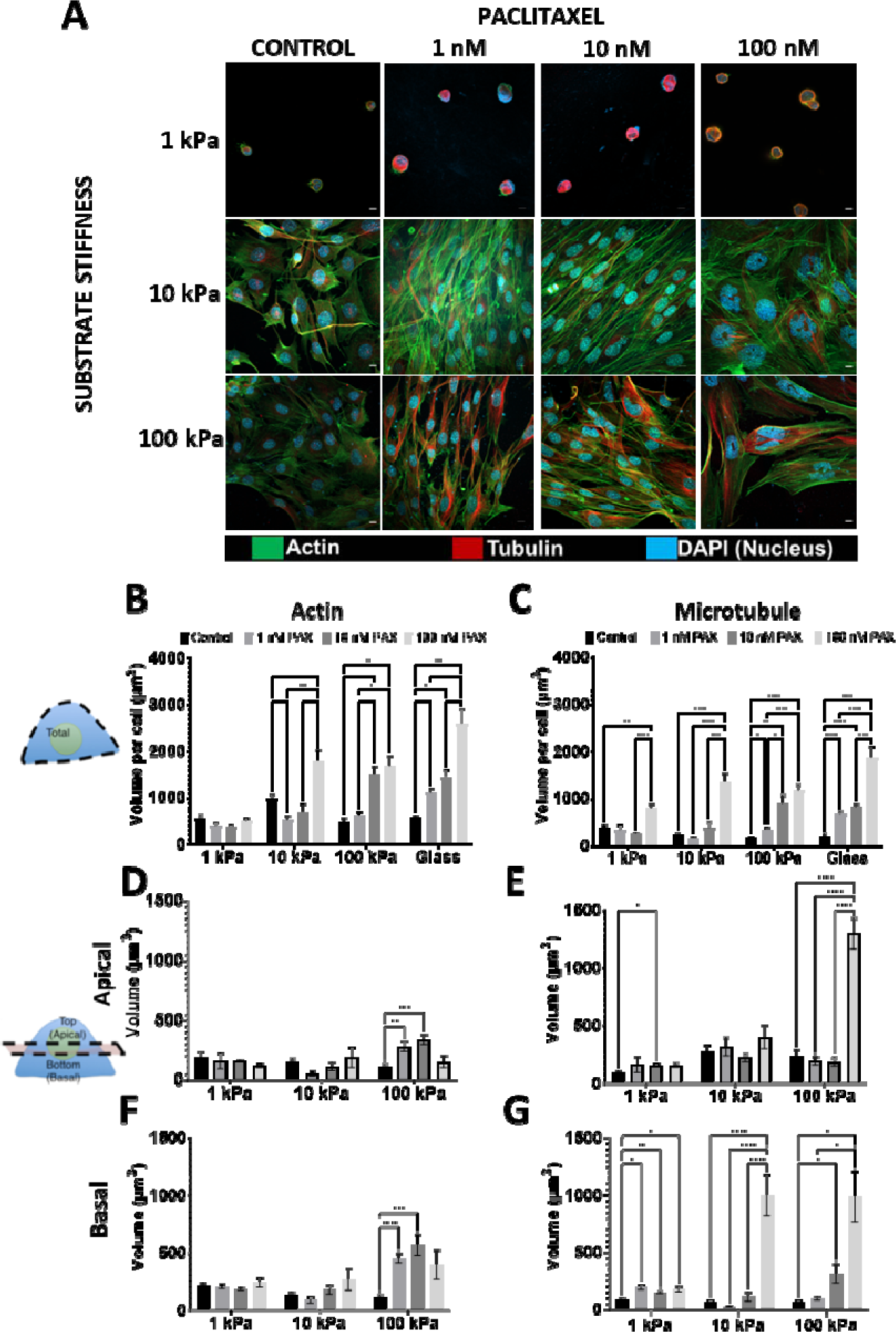
Changes in cytoskeletal spatial distribution provide a means for cells to adapt to substrate stiffness. (A) Culture on compliant substrates modulates cytoskeletal structure (scale bar 10 μm) upon stabilization of microtubule with PAX and the PAX concentration-dependent increase in (B) F-actin and (C) microtubule concentration per cell, apart from the 1 and 10 nM PAX on the softest (1 and 10 kPa) gel substrates. Segmenting the cell into apical and basal regions revealed fewer PAX-concentration dependent differences in respective volumes of F-actin (D and F) or microtubule (E and G). Taken as a whole, larger and more significant differences were observed in the amount of microtubule in the basal regions of the PAX exposed cells compared to controls.

#### Effect of more compliant substrates on F-actin alignment

Grossly visible changes in F-actin alignment occurred with both increasing substrate stiffness, from 1 to 100 kPa, as well as with exposure to increasing concentrations of PAX, except in cells seeded on the softest (1 kPa) gel substrates (Fig. 3A). In fact, cells cultured on 1 kPa gel and exposed to PAX showed **less** F-actin alignment than control cells not exposed to PAX (Fig. 3B). On 10 kPa and 100 kPa substrates, F-actin alignment of cells exposed to 1 nM PAX was lower than that of unexposed control cells (Fig. 3C, 3D); in contrast, on the same substrates, F-actin alignment of cells exposed to higher concentrations of PAX (10 and 100 nM) demonstrated respectively higher actin alignment than that of unexposed control cells (Fig. 3C, 3D), with respective differences slightly higher on the 100 kPa compared to the 10 kPa gel substrate. Actin alignment in cells cultured on 10 (Fig. 3C) and 100 kPa (Fig. 3D) gels showed similar alignment to cells cultured on glass (PART A Fig. 2A). Differences in the apical-basal alignment of the cytoskeleton were not distinguishable between PAX-exposed groups across the 1 (Fig. 3E), 10 (Fig. 3F) and 100 (Fig. 3G) kPa gel substrates. Hence, it appears that F-actin alignment is more strongly associated with mechanoadaptation to increasing stiffness substrates which are significantly **stiffer than the cells themselves**, i.e. 10 and 100 kPa gel substrates and glass, and is shifted slightly higher with inhibition of depolymerization via exposure to PAX at 10 and 100 nM.

**Figure 3.**
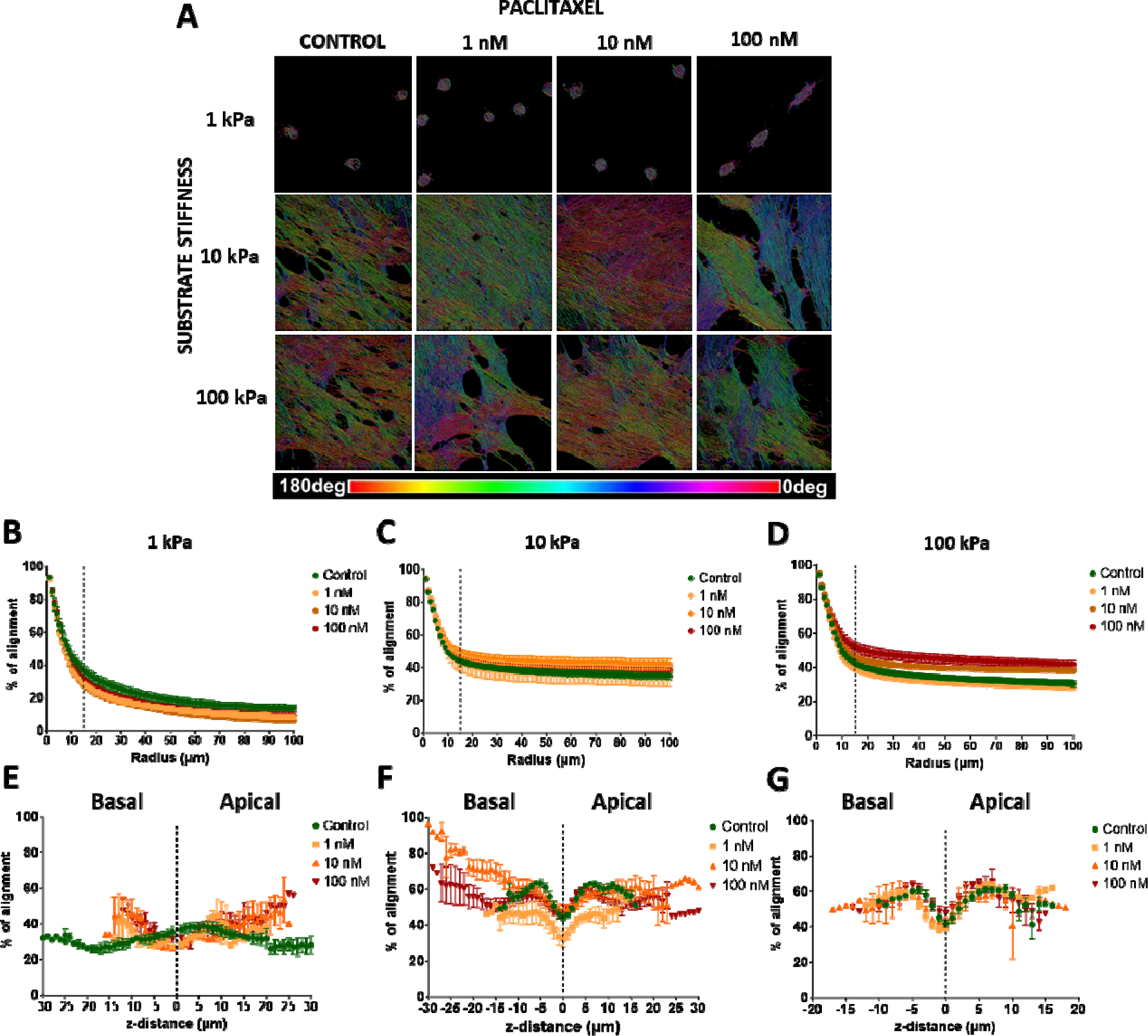
Actin filament alignment serves as a major change in phenotype during stem cell adaptation to substrate stiffness and stabilization of microtubules with PAX. (A) NoBS of actin reveals how substrate stiffness modulates the PAX-induced actin ordering. After 72h culture on (B) 1 and (C) 10 kPa gels, PAX-treated cells show a lower degree of actin alignment and a smaller difference between treated groups. (D) On 100 kPa gel, PAX-treated cells exhibit a higher degree of actin alignment than the control cells, although not as significant as those seeded on glass. The differences in apical and basal actin alignment on the softer gels, 1 kPa (E), 10 kPa (F), and 100 kPa (G), across the PAX exposed groups, are less distinct than those on glass. Nuclear z-thickness is used to determine the mid-plane of the cell (marked as dotted line at 0 μm) and to define the apical (top) and basal (bottom) regions. Although a clear distinction between apical and basal region can be observed with increasing substrate stiffness, there is no difference in the apical-basal actin alignment between the control and PAX-treated cells on soft substrates. Asterisk(s) represent significant differences at **** p < 0.0001, *** p < 0.001, ** p < 0.01, * p < 0.05, Tukey’s multiple correlation).

#### Effect of more compliant substrates on cells’ mechanical properties

Cells do not possess a pre-defined stiffness. Furthermore, increasing intrinsic cell stiffness and/or cytoskeleton-generated forces may drive cell mechanoadaptation over the 72h observation period to balance forces at the boundary of the cell to its local environment. Hence, we next probed the stiffness (via AFM) of cells cultured on gel substrates and correlated changes in stiffness to actin alignment (as described above). Compared to control cells seeded on glass, stiffness of cells on compliant substrates more closely matched the stiffness of their underlying substrate, and this adaptive cell stiffening increased with PAX exposure. Considering control cells cultured on substrates of increasing stiffness, on 10 kPa (Fig. 4A) and 100 kPa gels (Fig. 4B), control cells showed a mean respective Young’s Modulus of 8.08 kPa and 15.25 kPa; previously [Part A] we found that control cells on glass substrates (4 ± 2e+06 kPa) exhibited a mean Young’s Modulus of 3.3 kPa. PAX-exposed cells exhibited higher stiffness (Fig. 4C), both on 10 kPa and 100 kPa gel substrates where they exhibited higher respective mean Young’s Moduli of 10.9 kPa and 23.08 kPa, as well as on glass where they exhibited a mean Young’s Modulus of 6.4 kPa.

**Figure 4.**
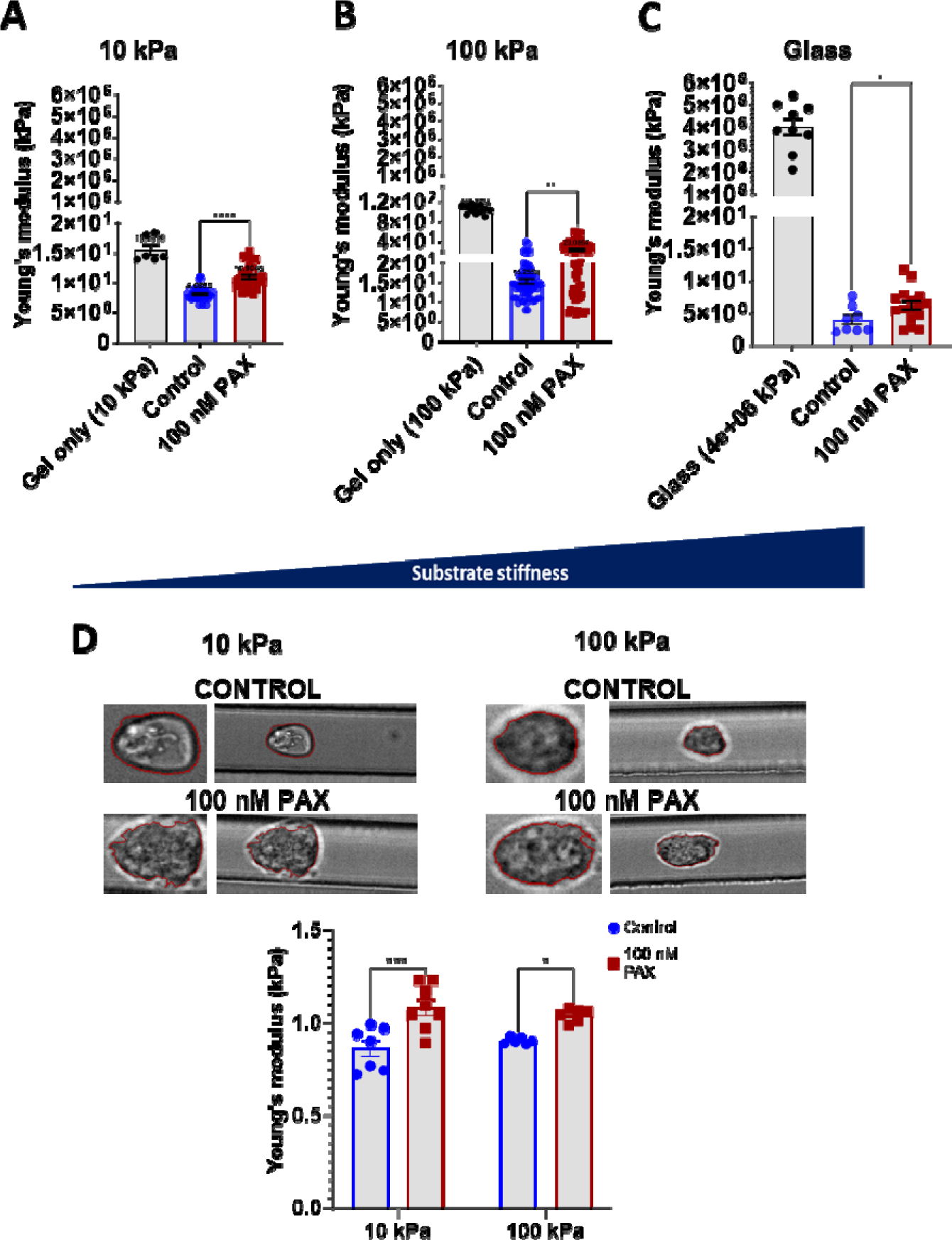
Young’s Moduli of cells cultured on substrates of increasing stiffness. Measurment via AFM revealed that on. (A) 10 kPa and (B) 100 kPa hydrogel, PAX-treated cells adapted more readily to match the stiffness of the gels than when cultured on glass (C). The differences in Young’s Moduli between the control and PAX-treated cells were more significant on the gels than on glass. (D) The significant stiffening effect of PAX exposure to cells was maintained even when the cells were trypsinized and the Young’s Moduli of the same cells in suspension were measured via deformability cytometry after culture on 10 kPa and 100 kPa gels and treated with PAX for 24h.

To probe mechanistically how this contrasting change in Young’s Modulus after PAX treatment could be modulated by underlying substrate stiffness, we harvested the cells after culture on gel substrates and exposure to PAX for 48h and measured the stiffness of the resulting suspended cells using deformability cytometry. The difference (increase) in Young’s Modulus between control and PAX-treated, suspended cells after culture on 10 kPa gel and 100 kPa gel, was as significant as the respective difference in Young’s Modul of suspended cells cultured on glass (Fig. 4D). The larger, irregular morphology of PAX-treated cells seeded on the gel could still be observed when they were flushed through the channel; however, they exhibited a smaller area (Fig. S2A) and volume (Fig. S2B) on 10 kPa gel than the comparable adherent cells. Cells were highly deformed in the cytometry channel (Fig. S2C). Similar differences (between adherent and nonadherent cells) were observed in cell area (Fig. S2D) and volume (Fig. S2), in cells seeded on 100 kPa gel substrates. The larger morphology attributable to PAX exposure also contributed to higher cell deformation within the channel (Fig. S2F).

We then examined how increasing substrate stiffness alone and in conjunction with local compression via seeding at increased density influences cells’ collective mechanoadaptation to substrate stiffness. First we measured the change from 24 to 72h in bulk modulus of cells seeded on substrates of increasing stiffness, *i.e.* proliferating from LD to HD over 72h. Cells exhibited bulk modulus stiffening resulting in a closer match to the stiffness of their respective substrates, with a mean Modulus of 7.7 kPa on the 10 kPa substrate and 22.1 kPa on 100 kPa substrate (Fig. 5A). Finally, exposure of cells to local compression by seeding at HD resulted in greater mechanoadaptation as measured by relative changes intrinsic cell stiffness than exposure to PAX (Fig. 5B). With PAX treatment, the mean bulk modulus of cells seeded at HD was higher than that of control cells, *i.e.* 13 kPa on the 10 kPa substrate and 28.3 kPa on 100 kPa substrate, a difference that is comparable to that of LD cells (Fig. 5B).

**Figure 5.**
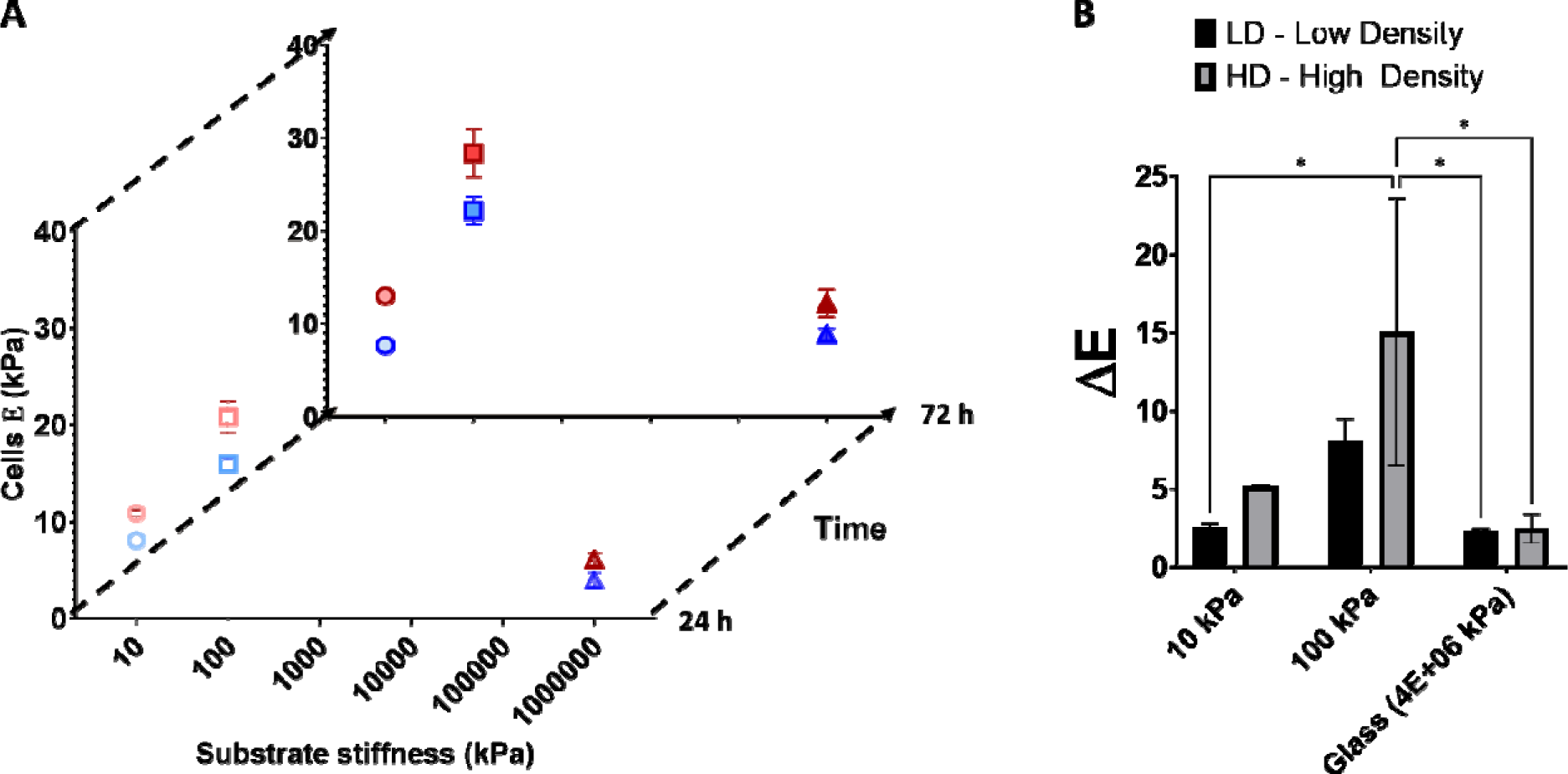
Differences in cells’ Young’s Moduli when cultured on substrate with increasing stiffness, demonstrating their adaptation in mechanical properties, and effect of seeding at higher initial densities, imparting local compression to cells. (A) Spatiotemporal changes of Young’s Moduli as they proliferated from low density (LD) at 24 h to high density (HD) at 72h and adapted to the different substrate stiffness during PAX treatment. (B) Changes in the Young’s Moduli (ΔE) of PAX exposed cells when seeded at LD or HD on different hydrogel substrates. Asterisk represents significant differences at * p < 0.05, Tukey’s multiple correlation).

### PAX-exposed stem cells adapt more readily to soft substrates via increasing actomyosin contractility

Despite the lower degree of actin alignment and its small difference after PAX treatment, the mean Young’s Moduli of stem cells cultured on soft substrates were higher than those cultured on glass, i.e. almost matched the Young’s Modulus of the gel. As stem cells show better capacity to adapt on softer substrates than on glass, with gels being more deformable or within stiffness ranges that cells can reciprocate with active pulling/tension, we then questioned the role of myosin II contractility in facilitating this adaptation. We probed for the expression of three myosin II isoforms that perform independent functions in maintaining cellular tension during adhesion^18^.

We found that the expression of Myosin IIA and IIB increases with increasing substrate stiffness, whereas the expression of Myosin IIC decreases (Fig. 6A). PAX treatment enhances the expression of the three Myosin II isoforms, i.e. Myosin IIA, IIB and IIC. Notably, the increase in Myosin IIA expression upon PAX treatment is higher on glass than on softer gels (Fig. 6B). In contrast, the increase in Myosin IIB and IIC expression is much larger on soft substrates than on glass (Fig. 6C and 6D).

**Figure 6.**
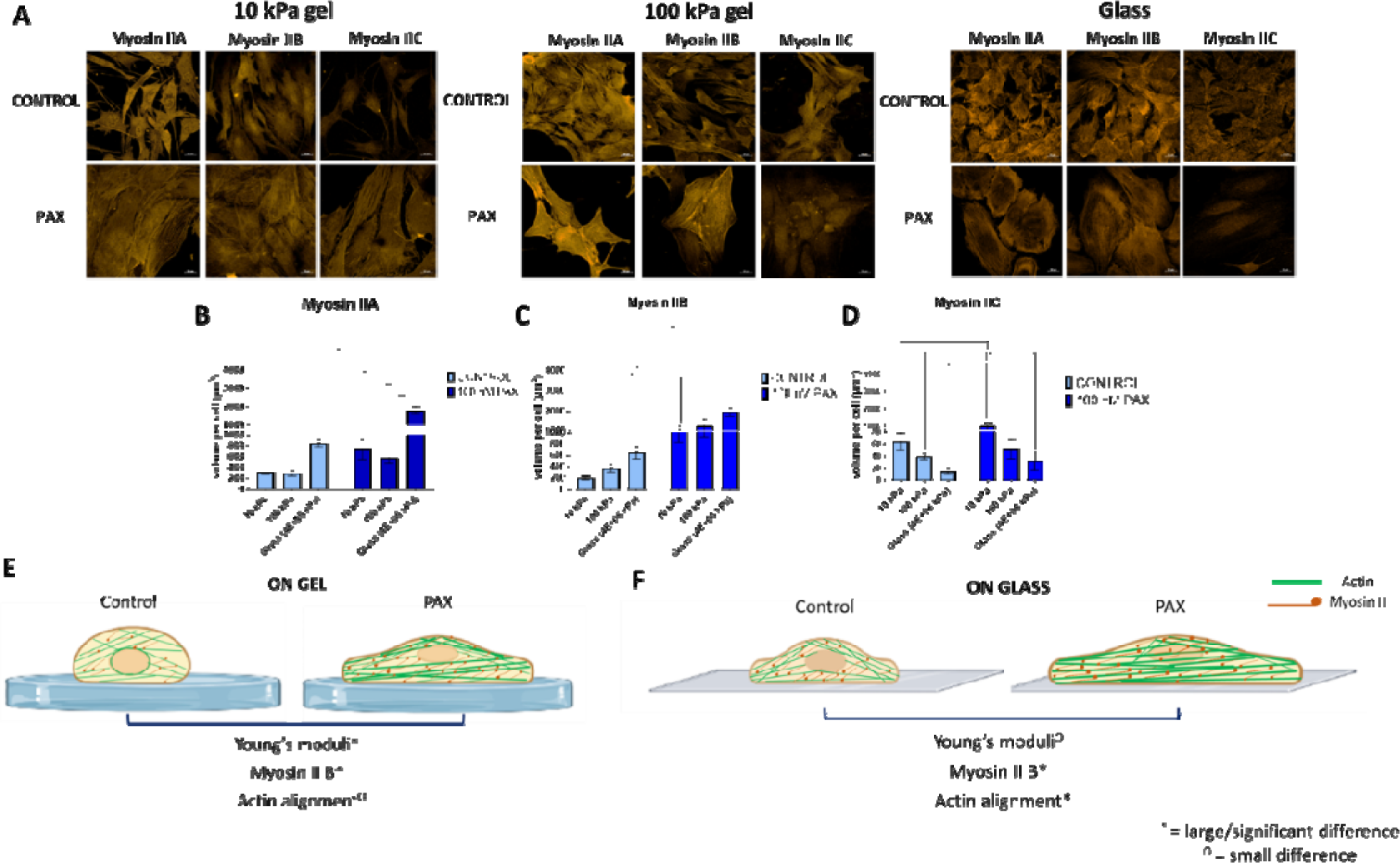
Stem cells adapt more readily on compliant substrates by means of their actomyosin contractility. PAX exposed stem cells exhibit significantly larger mean Young’s Moduli that those of control cells on compliant substrates, but not on glass, which corresponds with their significant increase in Myosin II expression (A), particularly Myosin II B. (B) Myosin II A expression as a function of substrate stiffness and PAX treatment. (C) Myosin II B expression shows significantly larger difference on softer substrates. (D) In contrast, Myosin II C expression decreases with increasing substrate stiffness but with larger difference with PAX treatment on softer substrates. Stem cell adaptation to substrate stiffness is b means of their actomyosin contractility that is more permissible on softer substrates (E) than on glass (F). Asterisk(s) indicate significant difference at * p< 0.05, ** < 0.01, *** < 0.001, and **** < 0.0001 (Tukey’s multiple correlation).

We summarize the experimental results regarding Myosin II contractility on stem cell adaptation to soft substrates in Fig. 6E and F. With relatively softer substrates, control cells exhibited more F-actin cross-links and short actin fibers oriented in random directions. With PAX exposure, a majority of stress fibers align within the cells; however, a greater number of short actin protrusions in random orientations and actin crosslinks between the aligned stress fibers were also observed. Taken as a whole, this not only contributes to lower global actin alignment when quantified with NoBS, but also accounts for the much larger difference in Young’s Modulus and Myosin IIB expression between the control and PAX-exposed groups. In contrast, for cells seeded on glass substrates, PAX induces long and thick, highly aligned actin stress fibers, resulting in higher Young’s Moduli and Myosin II B expression, although with less significant effects than for cells seeded on soft gels.

### Statistical correlations – testing correlations between independent and dependent variables

To test independent effects of increasing substrate compliance, exposure to PAX, seeding at increasing cell density, and decoupling of the cell from its native environment (suspension) we analyzed the correlation matrix between all dependent variables measured in this study (pooled data), indicating the level of interaction between cytoskeletal adaptation or remodeling parameters (Table 1). These dependent variables included cell volume, actin alignment, as well as actin and microtubule concentration (volume within cell). We defined significance in context of the experimental model system where exposure to PAX is known to stabilize polymerized microtubules by decreasing depolymerization; as such, the 0.332 correlation coefficient between per cell microtubule concentration and PAX concentration can be considered to be strongly correlated. By defining a physiologically relevant, strong correlation in this way, other calculated correlation values can be considered relative to that known strong correlation. Positive and negative, significant correlation coefficients corroborate results observed in each of the experimental studies described above and discussed in detail below.

**Table 1.**
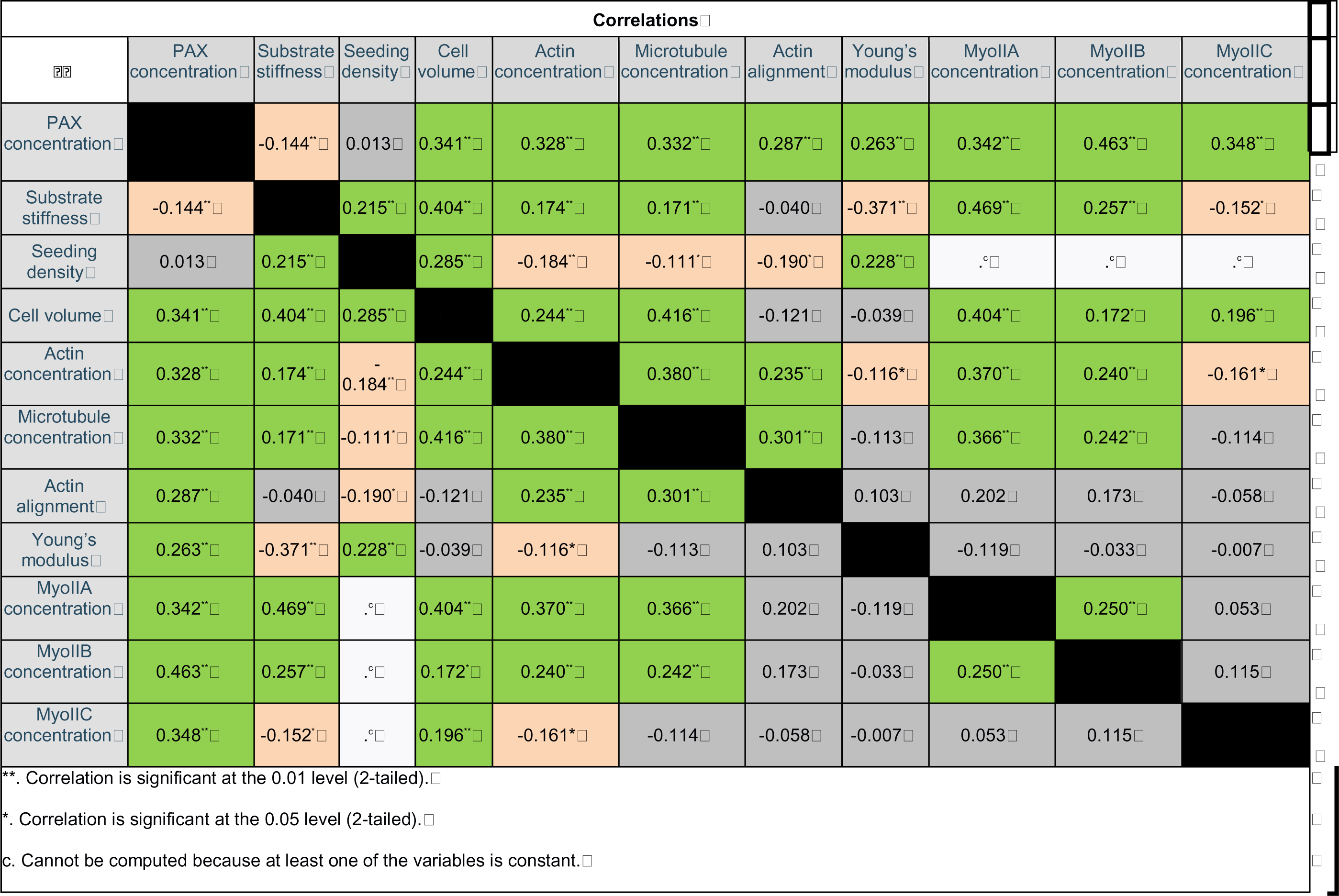
The correlation matrix showing the correlation coefficient between all independent variables (PAX exposure, seeding density, substrate stiffness) and outcome variables measured in this study (pooled data) and indicating the level of interaction between cytoskeletal adaptation or remodeling parameters: cell volume, actin alignment, actin and microtubule concentration. Green cells indicate a positive correlation (see details below), red cells indicate a negative correlation and gray cells indictes no correlation. Values indicate Pearson’s correlation coefficient. Note: In context of this experimental model system where exposure to PAX is known to stabilize polymerized microtubules by decreasing depolymerization, the 0.332 linear correlation coefficient between per cell microtubule concentration and PAX concentration can be considered as strongly correlated and other correlation values can be considered relative to that known correlation.

## Discussion

The current study (Part B) probed mechanoadaptation of mesenchymal stem cells in response to controlled biophysical cues known to induce local compression (seeding at increased density) and tension (seeding on compliant substrates), akin to mechanical testing of the most basic living and adapting unit of life. We exogenously perturbed the intrinsic capacity of MSCs to adapt via exposure to specific concentrations of PAX, which inhibited microtubule depolymerization, as developed in Part A of the study [Part A]. Yet stem cell mechanoadaptation to substrates of increasing stiffness was more pronounced than adaptation to exogenous cytoskeletal (tubulin) depolymerization inhibitors. Increasing PAX concentration up to 100 nM increased cell stiffness and volume, with associated emergent anisotropic cytoskeletal architectures. Probing of myosin II expression demonstrated that the MSCs adapted more readily on compliant substrates by virtue of their actomyosin contractility, i.e. active mechanoadaptation.

While stabilization of microtubules leads to the increase in actin length and alignment, providing a means to efficiently distribute larger forces within the cells, the degree of actin alignment is predominantly regulated by substrate (local environment) rigidity sensing. On glass, cells exhibit a clear PAX concentration-dependent stress fiber (SF) alignment, where at 100 nM PAX, a majority of SFs align in predominant directions. On compliant gels, this concentration-dependent increase in SF alignment is abolished and short, random F-actin cross-links are present along with aligned SFs, contributing to a lower alignment score. This is consistent with the reported regulation of fluid-like to solid-like actin behavior that depends mainly on substrate stiffness and not cell shape^10^. Furthermore, cells cultured on a circular shape pattern on a soft substrate possess high actin fluidity, though with increasing substrate stiffness actin remodels into higher order with unidirectional alignment while maintaining cell circular shape. Regardless of shape, stiff substrates prompt actin nematic behavior to exert larger traction force on opposite sides of cell periphery^10,11^. However, without shape confinement, cells would spread and initiate anisotropic focal adhesion points on the substrate, enabling a more effective force distribution.

In this study, SA/V, defining cell shape, increases with increasing PAX concentration in parallel with the increase of actin alignment on glass, whereas culture on gels seems to maintain SA/V across PAX concentration. This may be attributable to the fact that cells require a minimum area of spreading to survive^12^. Typical culture on glass or dishes provides unlimited adhesive substrate to the cells and hence when combined with PAX treatment compromises cell shape stability or isometric tension (pre-stress), which are otherwise maintained by the cell-cell and cell-extracellular matrix (ECM) interaction within physiological tissue^13^.

The cytoskeleton within the cell represents a system that maintains shape stability through adjustment/adaptation of pre-stress, the non-zero tensile stress of the actin filaments present when no external forces are acting on the cell boundary^14^. Microtubules are known to balance a portion of this tensile stress which serves as a means for cells to oppose changes in shape. However, when microtubule polymerization is disrupted (e.g., by colchicine) these stresses would shift to the substrate, resulting in an increase in traction force^15^, provided that the substrate is compliant. Hypothetically, when microtubules are stabilized with PAX, the stress would be instead borne by the cells causing actin SFs to grow linearly, unbranched, and thicker, which in turn would compromise cell shape as seen in the increasing SA/V. Particularly when the substrate is within the compliant range that cells can readily deform, dramatic changes in the F-actin observed in this study must occur to achieve minimal energy stored within the structure (force balance) and providing a mechanism whereby PAX treated cells could match the stiffness of the gel substrate. Cytoskeleton dynamics allows the unpredictable fluctuations of traction forces at the cell-substrate interface as a means to maintain tensional homeostasis^16^. Over time, the cytoskeleton maintains this fluctuation within a small range that could be matched with the mean traction force exerted onto the substrate^16^.

Simultaneously, cells have the capacity to adapt to their surrounding mechanical properties and tune their internal stress to match the stiffness of their underlying substrates^16,17^, which seems to be permissible within a certain range of substrate stiffness. On substrates of 0.5 kPa – 40 kPa, fibroblasts become stiffer as substrate stiffness increases before finally achieving a constant Young’s Modulus on 20 kPa substrates and above^8^. In this study, C3H/101/2 exhibit a mean Young’s Modulus of 3.3 kPa and about twice higher mean of Young’s Modulus of 6.4 kPa, upon PAX treatment, when probed on glass. Interestingly, cells reach mean Young’s moduli of 8.806 kPa and 15.25 kPa when probed on compliant 10 and 100 kPa gel substrates, respectively. PAX treatment increases cells’ Young’s Moduli to closely match the moduli of the compliant substrate, i.e. 10.9 kPa and 23.08 kPa on 10 and 100 kPa gels respectively. With microtubule disruption (PAX exposure), the energy stored in the compliant substrate is calculated as the work done by traction to deform the substrate, which in this case translates to the larger Young’s Moduli on compliant substrates than those on stiffer glass substrates.

Cell proliferation towards higher density represents the natural progression of cell growth and division, which influences how cells sense their local mechanical environment and dictate tissue morphogenesis^18,19^. When cells proliferate to higher density, the increase in cell bulk modulus upon PAX treatment was significantly larger on gel substrates than on glass. In the context of a multicellular construct, high density (HD) cells experience larger compression from the neighboring cells which balances the cells’ collective tensile forces. Particularly, tissues or multicellular constructs remodel extracellular matrix to achieve the balance of force exerted by the ECM on cells and forces generated by the cells themselves onto the ECM^20^. PAX treatment increases cell volume and microtubule pushing forces thus elevating the compression between cells. When cells were cultured on gel substrates, the excess energy from that pushing forces could be distributed and stored within the actin, microtubule and gel network, so that HD cells could readily achieve force balance – exhibiting stiffness that closely matches the gel substrates. HD cells proliferating to higher density are evidently more tensile on soft gels than on stiff gels^21^. On softer substrates HD cells in multiple 3D layers sense rigidity by dynamically adjusting their adhesion and tuning the low or high interactions with the substrate, allowing them to be more or less tensile^21^.

While it is counterintuitive that cells exhibit reduced stress fiber alignment on softer substrates despite their higher Young’s Moduli compared to stiff glass substrate, cells on compliant substrates exhibit more short actin structures with random orientation across the basal to apical regions and higher expression of Myosin II isoforms, indicative of contractility. In this study, we use exogenous PAX exposure to decipher the relationship between actin fiber rearrangements and cell stiffness adaptation, as a means to distribute stresses efficiently, particularly for cells cultured on gel substrates. Compliant substrates allow cells to exert larger traction force by means of randomly oriented and shorter F-actin, or via actin rearrangements and myosin contractility. In particular the larger increase in Myosin IIB and IIC expression on soft gels compared to stiff glass demonstrate the activity of those isoforms as endogenous stress-generation-dependent versus (over) purely substrate-stiffness-dependent. Given the role of Myosin IIB in bearing larger loads (higher duty ratio) and in polarization of traction forces^22^, PAX-stabilized microtubules might require a higher increase in IIB and IIC in conjunction with the increase in actin crosslinks. In contrast, Myosin IIA, as the major Myosin II isoform^22^, increases its expression with increasing substrate stiffness, and plays a critical role in generation of traction force and determination of cell polarity. Myosin IIA and IIB are reported to perform contrasting mechanical roles in cell spreading, whereby Myosin IIA retracts the lamellipodia extension mediated by Myosin IIB^23^. Hence, the difference in the increase of myosin isoforms’ expression, cellular mechanical properties, and contractility observed in this study during the modulation of endogenous stress by PAX, demonstrate the emergent adaptation to local environmental forces such as substrate rigidity and local compression. This emergent adaptation of F-actin and microtubules allows single cells and/or multi-cellular constructs to be more tensile, and to tune their adhesion to a range of substrate stiffnesses. With respect to development and healing, such emergent adaptation is important for cells to establish the structure-function relationship of the tissues they build^24^.

In summary, we demonstrate that by emulating stresses inherent to development, e.g. the local compression from increasing seeding density and local tension from compliant substrates, one can inform the capacity of cells (under perturbation by PAX) to achieve force balance via opposing shape changes and cytoskeletal reorganization (Supplementary Video 1). We also revealed the interactions between independent variables (environmental cues) including PAX concentration (inhibiting tubulin depolymerization), increasing seeding densities (local compression), and substrate stiffness (local tension) in influencing the degree of F-actin and microtubule spatial distribution and reorganization, as well as cell stiffness. The cells’ emergent adaptation to these local environmental cues provides information on the capacity of cells and the cytoskeleton to achieve tensional homeostasis in response to perturbation of cytoskeletal dynamics as well as changing mechanical and biophysical environmental cues intrinsic to tissue development in health and disease.

## Materials and methods

### Cell culture and Paclitaxel treatment

The C3H/10T1/2 murine embryonic stem cell line (CCL-226) was used as a model for primary embryonic mesenchymal stem cells and cultured in basal eagle medium (BME) with 1% penicillin-streptomycin, 1% L-glutamate, and 10% fetal bovine serum (FBS), per previously published protocols^7,25^. Cells were expanded and passaged to less than P15. For imaging, cells were seeded in glass-bottomed 24 well plates or 35 mm dishes coated with gel substrates of defined compliance (respective stiffness). For inducing local compression through seeding at increasing density^24–26^, cells were seeded at 5,000 cells/cm^2^ for low density (LD), 15,000 cells/cm^2^ for high density (HD) and 45,000 cells/cm^2^ for very high density (VHD).

Exogenous PAX exposure was carried out on the day following seeding. PAX solution was prepared from the stock of 5 mg/ml in Dimethyl Sulfoxide (DMSO) (5855 µM) in culture medium in a defined concentration range (1 – 100 nM). Medium was removed from the well plates and PAX solution was added to the wells. Control cells were treated with medium containing only DMSO. Cells were incubated until used for assays or imaging at determined time points (24, 48, and 72h).

### Polyacrylamide gel synthesis

Polyacrylamide gels, of 1, 10, and 100 kPa stiffness, were prepared as per our previous protocol^27,28^. Coverslips (18 mm) were first treated with 0.5% 3-aminopropyltrimethoxysilane solution, and then with 0.5% glutaraldehyde solution. Glass slides were made hydrophobic by treatment with RainX. A mixture of polyacrylamide was prepared by varying the ratio of acrylamide and bis-acrylamide monomers to achieve the desired stiffness. One mL of the polyacrylamide - acrylamide - bisacrylamide mixture was combined with 10 μl of 10% ammonium persulfate and 0.1 μl tetraethylmethylenediamine (TEMED). 20ul of the resulting mixture was pipetted onto the treated coverslips, after which the coverslips with gel were immediately turned upside down, facing the treated glass slide. The gels were left to polymerize for 10 minutes. The gels were then treated with 55% aqueous hydrazine hydrate for 2 hours, followed by washing with deionized water and glacial acetic acid for 1 hour each. Fibronectin was added into 3.5 mg/ml sodium periodate solution and was incubated for 30 minutes to oxidize the protein. 200 μl of the protein solution was pooled onto a polydimethylsiloxane (PDMS) stamp and incubated for 1 h. The stamps were air dried and then placed face down on to the PA gel surface. The stamp was lifted carefully, resulting in a gel with fibronectin surface coating, which was washed with deionized water and PBS and then sterilized under UV before being used as cell culture substrate.

### Cell and cytoskeleton imaging and analysis

Cells were labeled with Calcein AM and Hoechst stain for imaging of cell and nucleus structures. Actin was labelled using Actin Green^TM^ (LifeTech) with fluorophores that excite at 488nm. Microtubule cytoskeleton, as well as the three myosin II isoforms, Myo IIA, B, and C were labelled using antibodies-based incubation (Anti tubulin antibody (LifeTech Cat. No. A11126, 1:300 dilution), anti-Myosin IIA and anti-Myosin IIB (BioLegend, Poly19098 and Poly19099, 1:250 dilution) and anti-Myosin IIC antibody (Cell signalling, Cat. No. 3405, 1:300)) as described in the preceding study (Part A). The analysis of cell shape and volume, as well as cytoskeleton spatial distribution and actin fiber remodeling were quantified using custom MATLAB and Image J macro scripts as described in the preceding study [Part A].

### Atomic Force Microscopy (AFM)

Cells were seeded on to fibronectin coated polyacrylamide gels of various compliance at low density (LD), i.e. 5000 cells/cm^2^ and high density (HD), i.e. 15,000 cells/cm^2^ and treated with PAX for 24-72 h. Medium was replaced with serum free and phenol red free standard medium. Cells were taken for AFM measurement and analysis of cell Young’s Modulus following the protocols described in the preceding study [Part A].

### Deformability cytometry

Cells were seeded on to polyacrylamide gels on 25 mm coverslips in 6 well plates. Cells were treated the next day with 100 nM PAX or DMSO for control. After 48h, the cells were harvested and centrifuged, and the pellet was washed in PBS and resuspended in Cell Carrier buffer (Zellmechanik) for deformability cytometry analysis as described in the preceding study [Part A].

### Statistical analysis

Statistical analysis of experimental data was performed using Graphpad prism (La Jolla, CA) and SPSS (IBM). Significant differences in cell volume, stiffness, actin and microtubule concentration across PAX concentration, substrate stiffness and seeding densities were analyzed with Two-way ANOVA and Tukey’s multiple comparison test. A linear mixed model test was performed using the SPSS to investigate the effects of interacting variables on cells’ Young’s Modulus; this included a test of three-way interaction between independent variables: cell seeding density, substrate stiffness and PAX concentration, where data from each condition for each cell and repeat were pooled (Table S1). Bivariate Pearson’s correlation analysis between the dependent variables such as cell volume, actin and microtubule concentration, actin alignment, and Young’s Modulus, was performed to test how correlated their measures were from the three independent variables. The computed Pearson’s correlation coefficient with two-tailed test of significance was used to define the negative, positive correlation or no correlation (Table 1). As not all possible combinations of variables were tested in this study, two-way ANOVA was also performed to test the interaction between two independent variables in modulating cell volume, actin and microtubule concentration, and actin alignment (Table S2).

## Data Availability Statement

All data, documentation and code used in analysis is contained in the manuscript and Supplementary Material and can be provided upon request to the Corresponding Author.

## Acknowledgments

The authors would like to acknowledge the Katharina Gaus Light Microscopy Facility (KGLMF) and their staff for the continuous support in imaging resources and assistance in image processing and analysis. Special acknowledgement to Dr. Michael Carnell for developing the NoBS algorithm for the analysis of F-actin stress fibers alignment; Dr. Elvis Pandzic for assisting and training V.P. in custom-built MATLAB scripts for the analysis of cell shape, volume; and Dr. Celine Heu for assistance with the AFM; Dr. Chantal Kopecky for assistance in deformability cytometry. V.P. would like to thank the Scientia PhD Scholarship scheme from UNSW for support throughout her doctoral research studies. This work has been supported in part by grants from the National Health and Medical Research Council (MLKT, KAK) and the National Institute of Health (KAK).

