## Supplementary Information Figures and Legends for "Stem cell mechanoadaptation Part B - Microtubule stabilization and substrate compliance effects on cytoskeletal remodeling"

ORCID for KAK: 0000-0002-8963-9796

ORCID for MLKT: 0000-0002-3552-2525

**This PDF file includes:**

Figures S1 to S2

Tables S1 to S2

Legend for Supplementary Video 1

SI References

**Other supporting materials for this manuscript include the following:**

Animation SV1

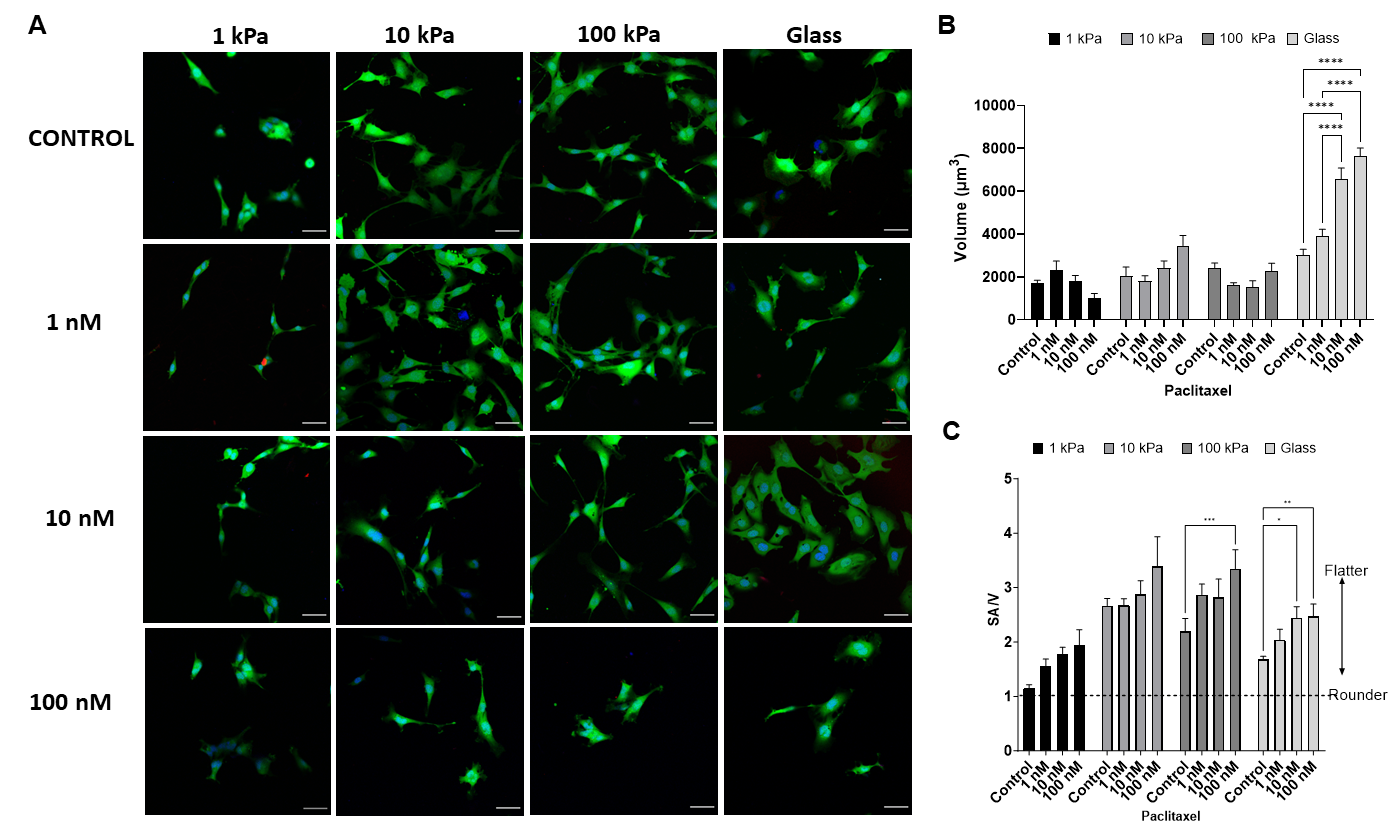

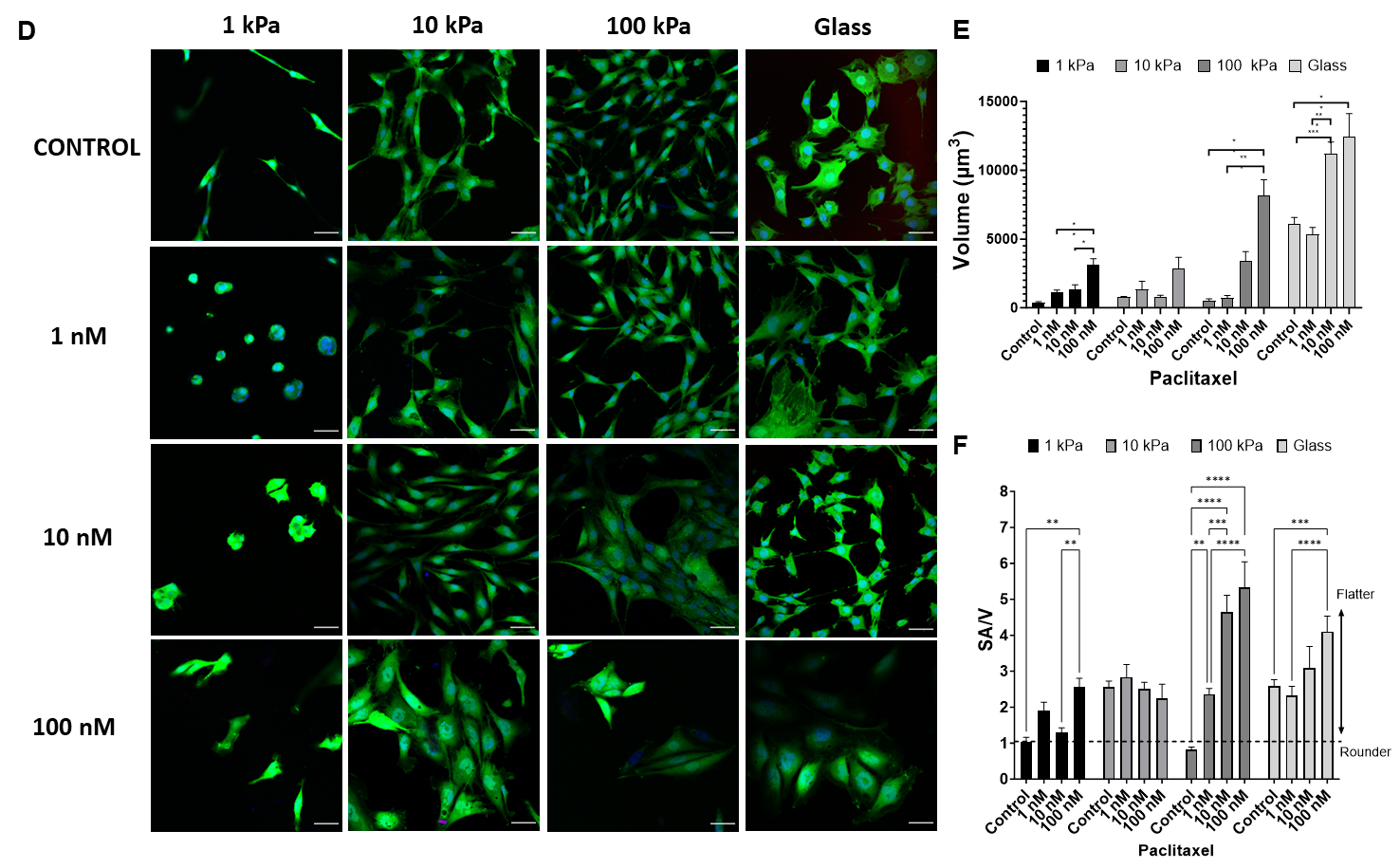

**Fig S1. Modulation of PAX-induced cell volume increase and shape changes upon introduction of compliant gel substrates**. (A) Substrate stiffness of 1, 10 and 100 kPa exert a profound influence in modulating PAX concentration-dependent cell volume increase (B) and shape changes (C) at 24. Culture on compliant substrates (D) and 48h PAX treatment were shown to modulate cell volume increase (E) and shape changes (F).

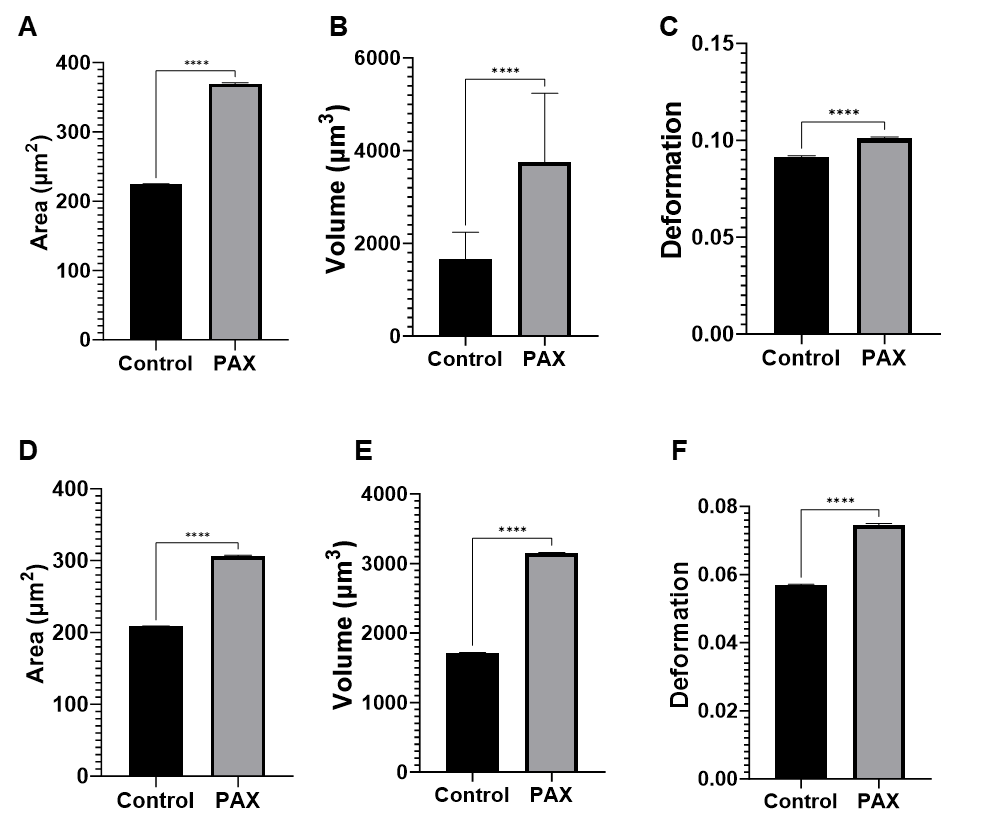

**Figure S2. PAX treatment consistently leads to significant increase in cell area and volume even when cells were previously cultured on compliant gels.** Measurement using deformability cytometry (Zellmechanik) revealed the increase in cellular area (A) and volume (B) after cells were cultured on 10 kPa and treated with PAX were significant. The increase in cell volume and size due to PAX treatment on 10 kPa (C) increased cell deformability. Culture on 100 kPa gel consistently maintained the increase in cell area (D) and volume (E) gels after treatment with PAX. Measurements were carried out for cells harvested in suspension. On 100 kPa gels (F), the increase in cell volume rendered cells more deformable when flushed through the deformability cytometry channel (30 µm width)

**Supplementary Animation 1, SV1. Force balances at cell boundaries modulate stem cells’ adaptation to their dynamic, local mechanical environment**. The force balance at the cell boundary is defined by the control volume (indicated by red dashed circle, with A – Area, and V – volume, and surface area of the cell) delineating the border between the outside of the cell (matrix and/or substrate in cultured cells, “out”), including the nucleus. The stress (σ) and strain (ε) relationships are delineated by theory of elasticity and plasticity, and theory of virtual work^1^. If the force balance (ΣF) is greater than zero (1), the cell experiences local compression. If the force balance (ΣF) is around zero (2), the cell is at steady state. If the balance (ΣF) is less than zero (3), the cell experiences local tension.

**Under each local mechanical condition (compression, steady state, tension), cellular adaptation ensues via either passive or active means.**

(1) Under local compression, one possible adaptive outcome is to compress to a smaller volume, either passively or through activation of ion channels, releasing intracellular fluid to the extracellular matrix. Alternatively, (active) cell division effectively increases the total cellular surface area to balance forces once steady state is reached. A further alternative is for the cytoskeleton to stiffen, effectively increasing the cell’s capacity to resist compression, and balancing forces to reach steady state.
(2) Under local compression, if forces are balanced at the cell boundary, no adaptation is required.
(3) In contrast, under local tension, expansion of cell volume, either passively or through activation of ion channels, provides a mechanism to balance forces. Based on results of the current experiments, this volume expansion leads to nuclear fragmentation due to tension transduced through the cell the nuclear envelope. A further adaptation alternative is cell division, which provides a means to increase total cell surface area and distribute the forces over a greater area. An additional alternative is for the cytoskeleton to soften, effectively increasing the cell’s capacity to expand under tension and balance forces.

**In different contexts, both *in vivo in situ* in the body**^2^ **as well as *in vitro* in cultured cells**^3^**, cells experience different force balances that are impacted by (A) seeding density and protocol**^3,4^**, (B) local compression, (C) local tension, e.g. with increasing substrate compliance, and (D) modulation of cytoskeletal stiffness, e.g. using microtubule stabilizer PAX.** In this study, each set of experiments probed A-D above.

(A) **Effects of substrate compliance versus seeding at density.**

Cells seeded at 5000 cells/cm2 and proliferated to higher density continue to divide contact inhibition occurs, *I.e.* when confluence is reached. Cells proliferated to target densities are more sensitive to changes in substrate compliance than cells seeded at target density, because the substrate exerts more influence on the basal boundary of the cells than surrounding cells exert when cells are seeded at density. When seeded on glass, which has very low compliance (4 x 10^6^ kPa), the cells are offloaded by the basal substrate. Seeding on a stiff hydrogel that is more compliant than glass (100 kPa) but less compliant than the cells themselves, the cells are offloaded, albeit to a lesser extent. Even when seeded on a 10 kPa hydrogel, the cells are somewhat off loaded along their basal surface. In contrast, when seeded on a hydrogel substrate similar in compliance to the cell itself (1 kPa), the seeded cells are no longer off loaded and they “round up”. Conversely, cells seeded at density are less sensitive to substrate compliance.

**(B) Local compression: effects of pressure difference at the cell membrane, tested**

**experimentally by seeding at increasing cell densities, which has been shown to exert local compression to cells**^3,4^**.**

Conceptually, in terms of the control volume (CV) of a single cell (red dashed line) within the

multicellular template created by seeding at density, if the stress inside the CV is less than that

outside of the CV, the cell experiences compression. Previous studies showed that this compression results in both a decrease in cell volume as well as a change in the shape of the nucleus (shape shift due to compression)^4^. Decreasing cell volume results in an effective stiffening of the cytoskeleton, both passively (increasing the density of the cytoskeleton via volume decrease) and potentially as an active adaptation via increased gene expression and cellular assembly. If the stress inside the cell CV is more than that outside of the CV, the cell experiences tension, resulting in a volume expansion and tension of the nucleus, which in this study was associated with nuclear fragmentation. Increasing cell volume results in an effective softening of the cytoskeleton, both passively (decreasing the density of the cytoskeleton via volume increase) and potentially as an active adaptation via decreased gene expression and cellular assembly.

**(C) Local tension: effects of increasing substrate compliance on force (stress) balance at the cell boundary (CV). The pressure differential is decreased (local tension) by increasing**

**substrate stiffness, or conversely by decreasing substrate compliance.** A more compliant cell is less stiff and less pressure resistant, while a stiffer cell is more resistant to pressure. A caveat in this conceptualization is that we do not know how far away from the cell boundary it can sense differences in compliance. Similar effects to (A) and (B) would be expected, with a working hypothesis that more compliant substrates mitigate effects of pressure differentials at the boundary of the cell (CV), shifting the cell toward relative tension. The impact of this effect would be greatest at lowest cell densities.

**(D) Effects of Paclitaxel (PAX), the microtubule stabilizing agent inhibiting depolymerization of tubulin.** In general, exposure to PAX increases cell volume and cell stiffness, effectively rendering the cell more compression resistant but potentially less tension resistant due to the observed fragmentation of the nucleus.

| **Type III Tests of Fixed Effects^a^** | | | | |
| --- | --- | --- | --- | --- |
| Source | Numerator df | Denominator df | F | Sig. |
| Intercept | 1 | 3.512 | 762.853 | 0.0000310917 |
| Paxconc | 1 | 370.494 | 79.777 | 0.0000000000 |
| substiff | 2 | 370.690 | 180.128 | 0.0000000000 |
| density | 1 | 370.527 | 43.530 | 0.0000000001 |
| Paxconc * substiff | 2 | 370.451 | 35.437 | 0.0000000000 |
| Paxconc * density | 1 | 370.246 | 14.726 | 0.0001461489 |
| substiff * density | 2 | 370.685 | 22.403 | 0.0000000007 |
| Paxconc * substiff * density | 2 | 370.584 | 15.935 | 0.0000002296 |
| a. Dependent Variable: youngsmod. | |  |  |  |

**Table S1. Tables of factors and interaction between three independent variables;** PAX concentration (PAX conc), substrate stiffness (substiff) and seeding density for cell stiffness and the cytoskeleton adaptation. The interactions effect on cell stiffness or Young’s moduli tested using linear mixed model (SPSS).

| **Interaction significance with P-value from Two-way ANOVA** | | | | |
| --- | --- | --- | --- | --- |
| **Factor** | **Cell volume** | **Actin concentration** | **Microtubule concentration** | **Actin alignment** |
| PAX concentration | Yes (<0.0001) | Yes (<0.0001) | Yes (<0.0001) | Yes (0.0003) |
| Substrate stiffness | Yes (0.0001) | Yes (<0.0001) | Yes (<0.0001) | Yes (<0.0001) |
| PAX concentration * Substrate stiffness | No (0.8488) | Yes (<0.0001) | Yes (<0.0001) | Yes (0.0266) |
| Seeding density | Yes (0.0015) | Yes (<0.0001) | Yes (<0.0001) | Yes (<0.0001) |
| PAX concentration * seeding density | Yes (<0.0001) | Yes (0.0063) | Yes (0.0003) | No (0.0541) |

**Table S2. Interactions** between two independent variables, PAX concentration and substrate stiffness or PAX concentration and seeding density, in influencing cell volume, actin or microtubule concentration, and alignment, tested with Two-way ANOVA.
